# Simulated drought with Polyethylene-Glycol (PEG) decreases above-ground performance and increases nodulation in the legume *Medicago lupulina*

**DOI:** 10.64898/2025.12.20.695366

**Authors:** Hana Cho, Emily Glasgow, Valmic Mukund, Julia A. Boyle, John R. Stinchcombe

## Abstract

Under climate change, understanding how plants and crops respond to drought is essential for basic research in ecology and evolution, and improving agricultural resilience. One common method of simulating drought in experimental conditions is by applying polyethylene glycol (PEG) to plants. We investigated drought growth responses in *Medicago lupulina* (black medic) using PEG to simulate drought stress. We grew *Medicago lupulina* plants inoculated with *Sinorhizobium meliloti* in Magenta boxes under controlled conditions and randomly assigned them to one of three treatments: a control, PEG applied to the bottom (PEG added to the bottom-watering container of a magenta box), or PEG applied from the top (PEG poured over the growth media). After 60 days, we measured true leaf number, nodule count, and below- and above-ground dry biomass. PEG treatments significantly reduced above-ground growth, including total biomass and leaf number, but unexpectedly increased nodulation. Our results suggest that while PEG effectively simulates drought stress on above-ground growth parameters, it may not accurately simulate drought effects on rhizobial symbiosis. PEG treatments had no effect on below-ground biomass, suggesting that increased nodulation is not a result of increased plant investment in below-ground growth under simulated drought. We hypothesize that PEG, as a persistent liquid that plants do not absorb, created conditions favorable for nodulation. Overall, these results highlight the importance of interpreting PEG-simulated drought experiments with caution when assessing mutualistic interactions.

## Description

With rapid and unpredictable climate change, the ability of crops to withstand drought is a growing concern (Juenger and Verslues 2023; Monteleone et al. 2023; Yuan et al. 2024). Water availability is an important factor that dictates plant evolution, ecological dynamics, and physiology (Juenger and Verslues 2023), and it is vital in agriculture, where water limitation reduces yields (Begna 2020; Moss et al. 2024). Therefore, as droughts become increasingly unpredictable and severe, understanding how plants respond and adapt to water stress is crucial for understanding plant population dynamics and evolution, and improving agricultural resilience (Begna 2020; Yuan et al. 2024). Here we investigated the drought responses of *Medicago lupulina* (Black medic) using simulated drought treatments with polyethylene glycol (PEG).

Manipulative experiments that simulate drought in the greenhouse, growth chamber, or laboratory are a useful tool for studying plant physiological and evolutionary responses to drought (Verslues et al. 2006), but they can be time and labor intensive. Common approaches include applying different watering regimes (Heschel et al. 2005; Puértolas et al. 2017), using shelters or devices to divert rainfall (Hoover et al. 2018; Boyle et al. 2024), conducting dry down experiments (Puértolas et al. 2017; Juenger and Verslues 2023; Boyle et al. 2025), or using Polyethylene glycol (PEG) (Verslues et al. 2006; Osmolovskaya et al. 2018). PEG is a widely used experimental tool because of its ease of use and because it creates osmotic stress without entering plant tissues (Burnett et al. 2005; Verslues et al. 2006). However, PEG-induced drought has been shown to decrease hypocotyl elongation, germination rate, ion uptake, and gene expression compared to those under real drought conditions (Burnett et al. 2005; Verslues et al. 2006; Osmolovskaya et al. 2018; Qi et al. 2023; Kylyshbayeva et al. 2024). Accordingly, there is a need to further characterize plant responses to PEG.

*Medicago lupulina* is an early successional, short-lived annual legume that plays an important ecological role through its symbiotic relationship with rhizobia, which facilitates nitrogen fixation and enhances soil moisture retention (Turkington and Cavers 1979). The species has a broad global distribution across North Africa, Europe, Asia, Australia, and North America, where it grows in dry grasslands, pastures, and along roadsides (Turkington and Cavers 1979; Clark 2008), and it is widely used as a forage and fodder crop (Heuzé et al. 2018). *Medicago lupulina* can grow at elevations up to 3,600 m in the Himalayas and tolerates a wide range of environmental conditions, including mean annual temperatures between 5.7°C and 22.5°C and annual precipitation levels from 310 mm to 1,710 mm (Duke 1981; Heuzé et al. 2018), suggesting it regularly experiences variable water availability.

We tested the effects of PEG-simulated drought on *Medicago lupulina* and its mutualistic interactions using three experimental treatments: a control, where plants received only water; a treatment where 150g L^-1^ 8000-PEG solution was poured over the growth media; and a treatment where the same solution was added to the bottom watering container of the magenta box (Figure 1a, see *Methods*; Burnett et al. 2005; Piwowarczyk et al. 2014; Castañeda and González 2021). We germinated seeds of *Medicago lupulina* and inoculated plants with mutualistic *Sinorhizobium meliloti* rhizobacteria. We applied nitrogen-free Jensen’s fertilizer (Boyle et al. 2021) after 26 days and ended the experiment after 60 days.

**Figure 1:**
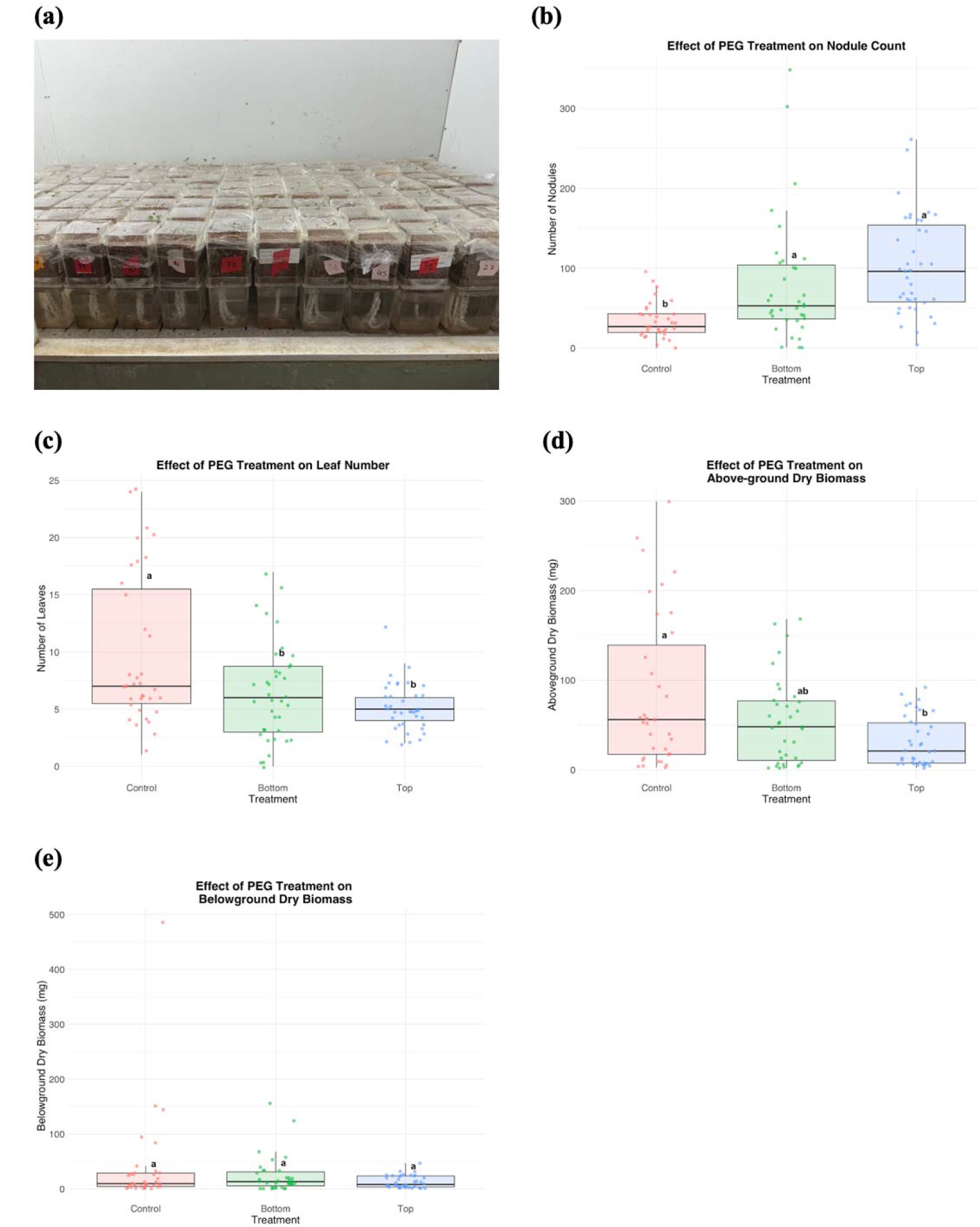
(a) Seed preparation and planting setup for Medicago lupulina. Individual seedlings were planted in Magenta boxes, covered with Saran Wrap and placed inside a growth chamber (25°C, 14-hour photoperiod). Effects of PEG treatments on Medicago lupulina growth. (b) Total nodule count, (c) number of leaves per plant across three treatments, (d) above-ground dry biomass (mg) and (e) below-ground dry biomass (mg): control, PEG applied from below (“Bottom”), and PEG applied from above (“Top”). Different letters indicate significant differences among treatments (Tukey HSD, p < 0.05).

Our results show that PEG treatment significantly reduced above-ground performance, specifically the number of leaves and above-ground biomass (Figure 1c and 1d). Control plants had significantly larger above-ground biomass than those top-watered with PEG solution, while those bottom-watered with PEG were not significantly different from controls or top-watering PEG-treated plants (Figure 1e). These data suggest that PEG can effectively simulate drought stress for above-ground growth metrics. However, we also detected a significant increase in nodule number in response to PEG application (Figure 1b). The usual effect of drought on nodulation is nodule senescence and decreased nodulation (e.g., Arrese-Igor et al. 2011; Iqbal et al. 2022; Istanbul et al. 2022; Melo et al. 2025; Ruiz□Lozano et al. 2001; for a phylogenetic meta-analysis, including species of *Medicago*, see Iqbal et al. 2022). Mhadhbi et al. (2009) showed decreased nodule number in *Medicago truncatula* in response to osmotic stress from mannitol treatments, while Dhanushkodi et al. (2018) showed that reduced soil water content in the *Medicago truncatula* led to an ∼50% reduction in mean nodule number relative to well-watered controls. Our findings of significantly increased nodulation in both PEG treatments relative to the control are thus in contrast with the prevailing reports in the literature for legumes, phylogenetic meta-analyses, and experimental results using simulated and actual drought in congeneric *Medicago* species. We originally hypothesized that plants might compensate for simulated drought by investing more in below-ground growth, which could explain higher nodule counts (Koziol et al. 2012; Sofi et al. 2018; Lumactud et al. 2023). However, our measurements of dry belowground biomass showed no significant differences among treatments (Figure 1e), which does not support this explanation. Multiple alternative hypotheses could explain the finding of increased nodulation in response to PEG. It could be that PEG solution, a persistent liquid that the plants could not absorb, was favorable for rhizobia mobility and/or infection, resulting in increased nodulation. Alternatively, PEG may affect plant production of flavonoids (Sarmadi et al. 2019), a key step in the nodulation process (Subramanian et al. 2007), or the dispersal of flavonoids away from the roots, both of which could have led to increased nodulation. In addition, osmotic stress has been shown to increase the expression of Nod genes in other species of *Sinorhizobium* (Fuentes-Romero et al. 2023), which if it occurred in this experiment, could also have led to increased nodulation. Regardless of the mechanism, our results suggest caution in the use of PEG to simulate the effect of drought on plant-rhizobia mutualisms.

## Methods

### Seed Preparation

We scarified a total of 150 *M. lupulina* seeds, sterilized them in 100% ethanol, and imbibed them in distilled water. The seeds were placed on 1% agar plates, wrapped in aluminium foil, and stored at 4 °C for eight days (Simonsen and Stinchcombe 2014; Boyle et al. 2021). We then moved them to room temperature (∼24 °C) for 16 hours to allow radicle emergence (Simonsen and Stinchcombe 2014; Boyle et al. 2021). Germinated seeds were planted individually in Magenta boxes. Each box consisted of a top chamber filled with sterilized Turface and a cotton string wick extending through a hole at the bottom to draw water from a lower chamber. The setup was covered with Saran Wrap in an attempt to minimize sources of dehydration other than those caused by PEG; when plants grew tall enough, we poked a small hole in the Saran Wrap and gently pulled stems through. We placed magenta boxes in a growth chamber with a 14-hour photoperiod at 25°C; we cultured the bacterial strain *Sinorhizobium meliloti* WSM1022 following Simonsen and Stinchcombe (2014), and applied 1 mL of inoculum the day after planting (Figure 1d). After 26 days of growth, we applied 1 mL of nitrogen-free Jensen’s fertilizer.

### PEG Treatments

We prepared 150 g L□^1^ polyethylene glycol (PEG 8000) solution to simulate drought. Plants were randomly assigned to one of three treatments: a control with 150 mL of sterilized water added to the bottom chamber, a “Bottom” treatment where 150 mL of PEG solution was added to the bottom chamber, and a “Top” treatment where 150 mL of PEG solution was applied to the soil surface. We included both the top and bottom treatments because it was unclear how the PEG solution would interact with the wicks used in the Magenta boxes; the top treatment ensured that plants and their roots would be exposed to PEG in the growth media. We applied the treatments 30 days after planting, with no further water or PEG solution supplied for the remainder of the experiment.

### Measurements

We counted the number of true leaves 40 days after they were planted. On day 50, we destructively harvested four plants to visually confirm nodule formation; these samples were excluded from further analyses. Because of these destructive samples, and some germinants failing to establish, final sample sizes were 43 controls, 44 in the bottom PEG treatment, and 43 in the top PEG treatment. We took measurements 60 days after planting, including true leaf and nodule counts, and dried above-ground dry biomass. We initially stored roots in a lab fridge, then oven-dried and weighed them to obtain below-ground dry biomass.

### Statistical Analysis

We conducted the analysis in R (v4.5.1; R Core Team 2025) using base R functions and the *tidyverse, ggplot2*, and *multcompView* packages for data processing and visualization (Graves et al. 2024; Wickham 2019; Wickham et al. 2019). For below-ground biomass, a single data point was 8.8 standard deviations from the mean and was removed as an outlier. We used a one-way analysis of variance (ANOVA) function, *aov ()*, to evaluate the effect of the different PEG treatments on leaf number, nodule number, and below- and above-ground dry biomass. When ANOVA results indicated significant treatment effects (p < 0.05), we conducted pairwise comparisons among treatments using Tukey’s Honestly Significant Difference (HSD) tests. Significance groupings were extracted using the *multcompLetters4()* function and we created a box plot with overlaid jittered data points to display individual variation. Raw data and R code for this work are archived at Zenodo (Cho et al. 2026).

## Acknowledgements

We thank the EEB horticultural staff for advice and support.

